# Maternal Western-style diet programs skeletal muscle gene expression in lean adolescent Japanese macaque offspring

**DOI:** 10.1101/2024.05.17.594191

**Authors:** Emily A. Beck, Byron Hetrick, Luis Nassar, Douglas W. Turnbull, Tyler A. Dean, Maureen Gannon, Kjersti M. Aagaard, Stephanie R. Wesolowski, Jacob E. Friedman, Paul Kievit, Carrie E. McCurdy

**Affiliations:** Data Science, University of Oregon, Eugene OR, USA; Department of Molecular Biosciences, University of Kansas, Lawrence, KS, USA; Department of Human Physiology, University of Oregon, Eugene OR, USA; Phil & Penny Knight Campus for Accelerating Scientific Impact, University of Oregon, Eugene OR, USA; Genomics and Cell Characterization Core Facility, University of Oregon, Eugene OR, USA; Division of Cardiometabolic Health, Oregon Health Science University, Oregon National Primate Research Center, Beaverton, OR, USA; Department of Medicine, Division of Diabetes, Endocrinology, and Metabolism, Vanderbilt University Medical Center, Nashville, TN USA; Department of Obstetrics and Gynecology, Division of Maternal-Fetal Medicine, Baylor College of Medicine & Texas Children’s Hospital, Houston, TX, USA; Department of Pediatrics, Perinatal Research Center, University of Colorado, Aurora, CO, USA; Harold Hamm Diabetes Center, University of Oklahoma Health Sciences Center, Oklahoma City, OK, USA

**Keywords:** metabolic programming, maternal diet, offspring, skeletal muscle, gene expression, maternal obesity

## Abstract

Early-life exposure to maternal obesity or a maternal calorically dense Western-style diet (WSD) is strongly associated with a greater risk of metabolic diseases in offspring, most notably insulin resistance and metabolic dysfunction-associated steatotic liver disease (MASLD). Prior studies in our well-characterized Japanese macaque model demonstrated that offspring of dams fed a WSD, even when weaned onto a control (CTR) diet, had reductions in skeletal muscle mitochondrial metabolism and increased skeletal muscle insulin resistance compared to offspring of dams on CTR diet. In the current study, we employed a nested design to test for differences in gene expression in skeletal muscle from lean 3-year-old adolescent offspring from dams fed a maternal WSD in both the presence and absence of maternal obesity or lean dams fed a CTR diet. We included offspring weaned to both a WSD or CTR diet to further account for differences in response to post-weaning diet and interaction effects between diets. Overall, we found that a maternal WSD fed to dams during pregnancy and lactation was the principal driver of differential gene expression (DEG) in offspring muscle at this time point. We identified key gene pathways important in insulin signaling including PI3K-Akt and MAP-kinase, regulation of muscle regeneration, and transcription-translation feedback loops, in both male and female offspring. Muscle DEG showed no measurable difference between offspring of obese dams on WSD compared to those of lean dams fed WSD. A post-weaning WSD effected offspring transcription only in individuals from the maternal CTR diet group but not in maternal WSD group. Collectively, we identify that maternal diet composition has a significant and lasting impact on offspring muscle transcriptome and influences later transcriptional response to WSD in muscle, which may underlie the increased metabolic disease risk in offspring.

## INTRODUCTION

The risk for the development of obesity and cardiometabolic disease, including type 2 diabetes in youth is increased by in utero exposure to maternal malnutrition or, paradoxically, a maternal caloric dense and high fat diet ^1–4^. As two thirds of women in the U.S. fall within or above the overweight BMI category^5^ and with many individuals having limited access to optimal nutrition^6–8^, the consequences and root sources of these developmental exposure(s) represent a substantial risk to the health of our population. Importantly, the pathologic processes that underlie diabetes, insulin resistance and beta-cell dysfunction, progress more rapidly in youth with type 2 diabetes (T2D) than in adults with the disease^9,10^. These factors, and a poorer response to treatment, result in overall worse glycemic control and in an increased risk of early diabetes-related complications including hypertension, microvascular-related diseases, and dyslipidemia in youth with T2D^11^. Still, the genetic and molecular mechanisms linking early life exposures to the increased incidence of T2D later in life remain a critical gap in knowledge.

Dysregulation of skeletal muscle insulin-signaling and metabolism is a hallmark of insulin resistance and an underlying cause of cardiometabolic diseases and the progression of age-associated dysfunction^12^. Prior work in our Japanese macaque model of Western-style diet (WSD)-induced maternal obesity identified metabolic dysregulation in multiple tissues of fetal, juvenile, and adolescent offspring,^13–18^ including reduced insulin sensitivity in skeletal muscle^19^ and decreased oxidative metabolism in fetal muscle^20^. Adaptations to a maternal WSD persisted in skeletal muscle in juvenile and adolescent macaque offspring^21^, and in rodents, the insulin resistant phenotype persisted for multiple offspring generations^22^, even when offspring are switched to a healthy diet. The persistence of these phenotypes in offspring exposed to a maternal WSD during pregnancy suggests persistent and durable molecular reprogramming and modifications to the transcriptome^23^.

Given the role of skeletal muscle in regulating peripheral insulin resistance during adolescence, we sought to determine the impact of maternal WSD feeding during pregnancy and lactation on programming of transcription along well-characterized pathways. Given the potential variation in adiposity deposition amongst males and females during these key lifespan intervals leading up to reproductive competence, we further sought to explore sex as a biologic factor using a nested experimental design. Specifically, in the current study, we aimed to investigate the effects of maternal WSD vs. maternal obesity plus a WSD (OB) vs. post weaning WSD. To this end, we compared effects of maternal (M) and/or post-weaning (PW) WSD feeding, compared to a standard control diet (CTR), on skeletal muscle gene expression in early adolescent 3-year-old Japanese macaques. This strategy resulted in 5 experimental treatment groups: Offspring from lean dams fed a M-CTR that were fed a PW-CTR (Group 1; CTR/CTR) or a PW-WSD (Group 2; CTR/WSD) and offspring from lean dams fed a M-WSD that were then fed a PW-CTR (Group 3; WSD/CTR) or a PW-WSD (Group 4; WSD/WSD) (**Figure 1A-C**), and offspring from M-OB weaned to a PW-CTR (Group 5; obWSD/CTR). All experimental groups consisted of both male and female offspring. (**Figure 1B**). Based on our prior work, we hypothesized that pathways related to skeletal muscle metabolism, growth and insulin signaling would be differentially expressed in offspring exposed to maternal or post-weaning WSD, when compared to those maintained on a CTR diet. Additionally, we anticipated male offspring to have less favorable transcriptional adaptations to maternal WSD than their female counterparts given reports of worse metabolic phenotypes in adult male offspring^24^.

**Figure 1.**
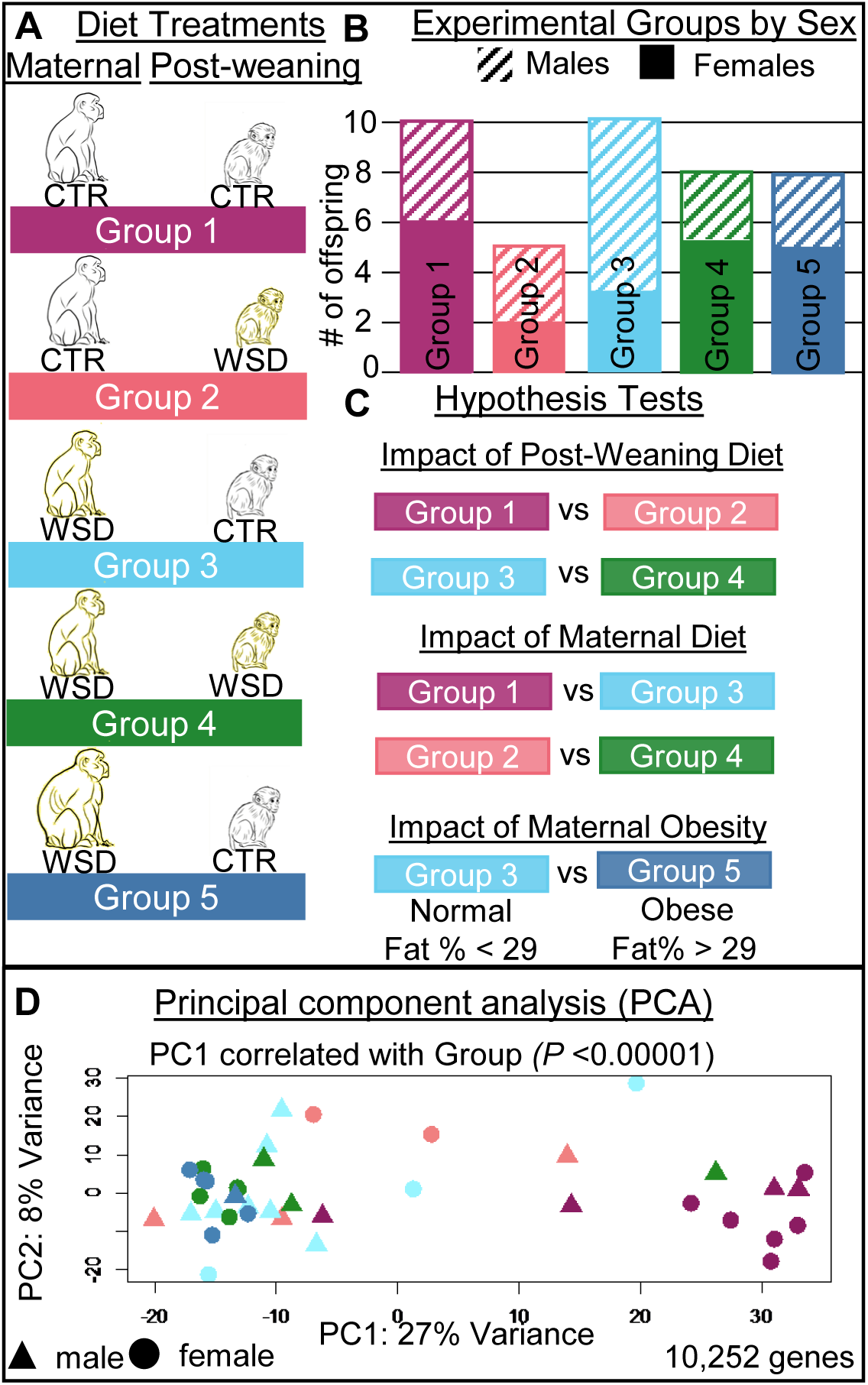
Experimental Design and Hypotheses Tested. Schematic representation of experimental group composition, group metadata, hypotheses tested, and visualization of total variance in transcriptional profiles. Color-coding of groups is consistent throughout the manuscript. Yellow shading of the macaque indicates western-style diet (WSD), black indicates a control diet (CTR). **(A)** Composition of the five experimental groups in this study. Group 1 (purple): offspring on a CTR diet with lean dams fed a CTR diet. Group 2 (pink): offspring fed a WSD with lean dams fed a CTR diet. Group 3 (light blue): offspring fed a CTR diet with lean dams fed a WSD. Group 4 (green): offspring fed a WSD with lean dams fed a WSD. Group 5 (dark blue): offspring fed a CTR diet with obese dams fed a WSD. **(B)** Stacked bar chart representing total sample size in each group and sex composition of each group. Hatched lines indicate number of males, solid fill indicates females. **(C)** Groups used for each hypothesis test. **(D)** PCA of all 39 offspring included in the study.

## RESULTS

### Offspring gene expression is significantly different based on experimental group

To identify differences between skeletal muscle transcriptional profiles, we implemented a Principal Component Analysis (PCA) combining all 40 samples across the five experimental groups (**Figure S1**). We identified a clear outlier within WSD/CTR (Group 5). In assessing read count distributions, we did not identify any difference in coverage for the outlier (**Figure S2**) and remove it from future analyses – only impacting tests of M-obesity (**Figure 1D**). Removing this outlier resulted in retainment of only a single male in Group 5 and therefore sex was not considered when assessing impacts of M-obesity on offspring gene expression. We then assessed relatedness of transcriptional profiles of the remaining 39 samples in PC space. After removal of the outlier, we observed a significant correlation between experimental group assignment and PC1 (*P* < 0.00001) with PC1 accounting for 27% of the variance observed (**Figure 1D**). As groups are parsed by three main variables (PW-diet, M-diet, and M-obesity) significant association of PC1 with experimental group warranted further exploration into the causal variable(s) for this partition of experimental groups.

### Maternal obesity does not significantly impact offspring gene expression

To test for impacts of M-obesity on offspring gene expression, we compared individuals from M-WSD and PW-CTR groups whose mothers were obese vs lean (Group 3 vs Group 5). Given the removal of an outlier from Group 5, we only retained a single male offspring from an obese dam and therefore could only test for effects of M-obesity with sexes combined (**Figure 2A**). We performed hierarchical clustering of the top 1000 differentially expressed genes which did not reveal any clustering by group (**Figure 2B**). The lack of clustering was also observed using PCA where there was no significant correlation of group with any PC (*P >* 0.05) (**Figure 2C**). We then fit the data to a linear model accounting for maternal obesity and identified only a single significantly upregulated gene (**Figure 2D**). These data suggest no significant effect of M-obesity on offspring skeletal muscle gene expression.

**Figure 2.**
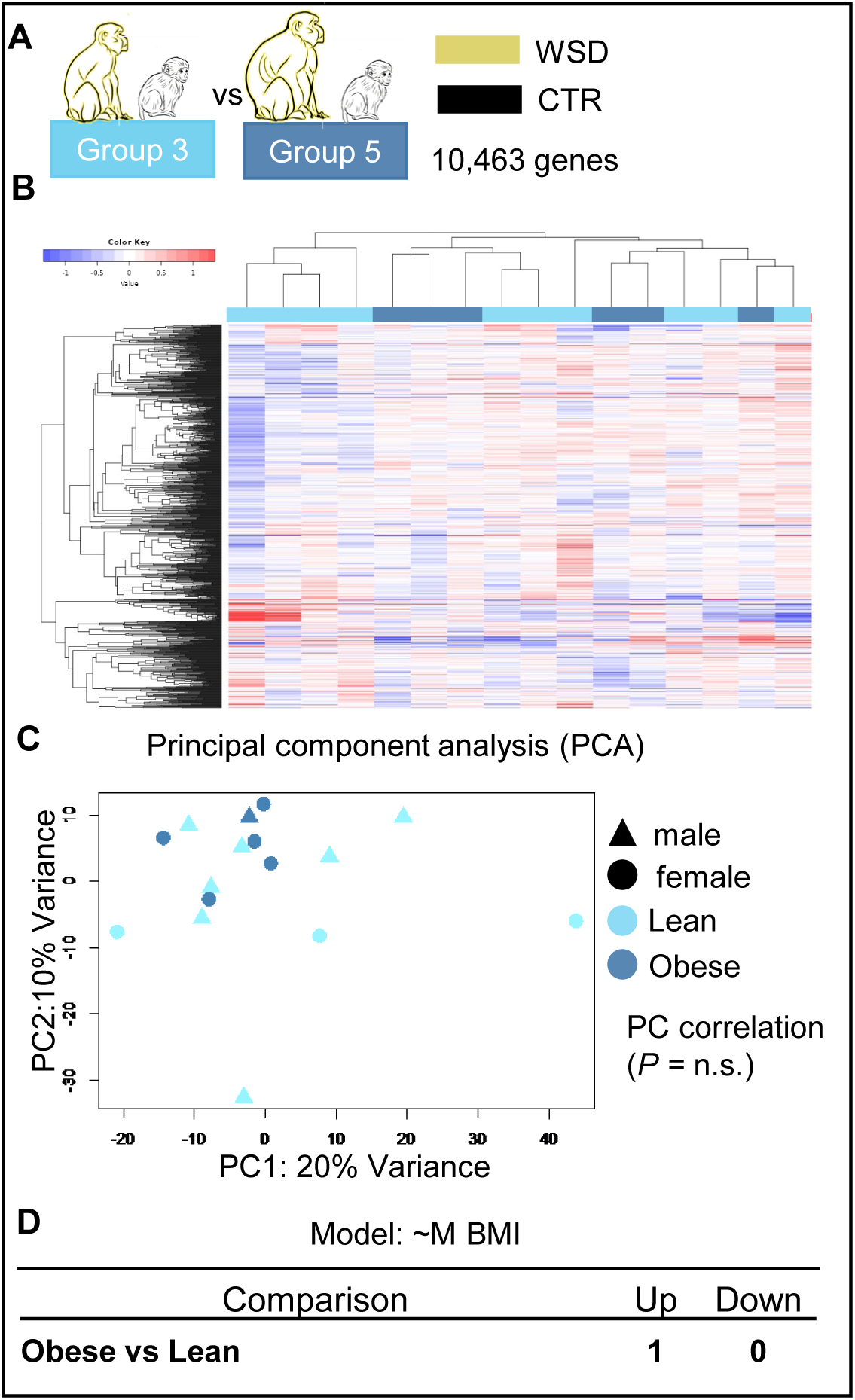
Impact of maternal obesity on offspring gene expression. Colors indicate experimental group with triangles representing males and circles representing females. **(A)** Group composition and the number of genes retained post-filtering. **(B)** Hierarchical clustering of the top 1,000 differentially expressed genes. **(C)** PCA of filtered transcriptional profiles. (**D)** Linear models and number of significantly differentially expressed genes by maternal obesity.

### Maternal WSD feeding significantly impacts gene expression

To assess the impact of M-diet on offspring skeletal muscle gene expression, we performed three comparisons. First, we compared PW-CTR groups exposed to either M-WSD or M-CTR diet (Group 1 vs. Group 3) (**Figure S3A**). As we did not observe any impact of M-obesity on offspring gene expression, we performed a third comparison combining WSD/CTR groups (lean and obese mothers) to be compared to CTR/CTR (Group 1 vs Group 3 & 5) (**Figure 3A**). Combining Groups 3 & 5 allowed us to increase our sample sizes and account for offspring sex. We performed hierarchical clustering of the top 1,000 differentially expressed genes – regardless of significance – in comparison to assess partitioning of individuals. We observed clear separate grouping of individuals from CTR/CTR vs. Lean WSD/CTR (**Figure S3B**). This pattern was consistent across comparisons including and excluding obese mothers (**Figure 3B**). PCA of each comparison confirmed these findings such that PC1 significantly correlated with group assignments in the Group 1 vs. Group 3 comparison and Group 1 vs Groups 3 & 5 combined comparison (*P* = 0.0001; *P* < 0.00001 respectively) with PC1 accounting for 32% of the observed variance (**Figure S3C; Figure 3C**). When combining Groups 3 and 5, we observed a significant correlation of read count by group (*P* = 0.042). When fitting our data to a linear model accounting for M-diet, we identified 272 significantly upregulated and 620 significantly downregulated genes in PW-CTR groups comparing M-WSD to M-CTR diet (i.e. Group 1 vs. Group 3) (**Figure S3D**). The number of impacted genes increased with inclusion of offspring from obese mothers to 338 significantly upregulated and 682 significantly downregulated genes (**Figure 3D**).

**Figure 3.**
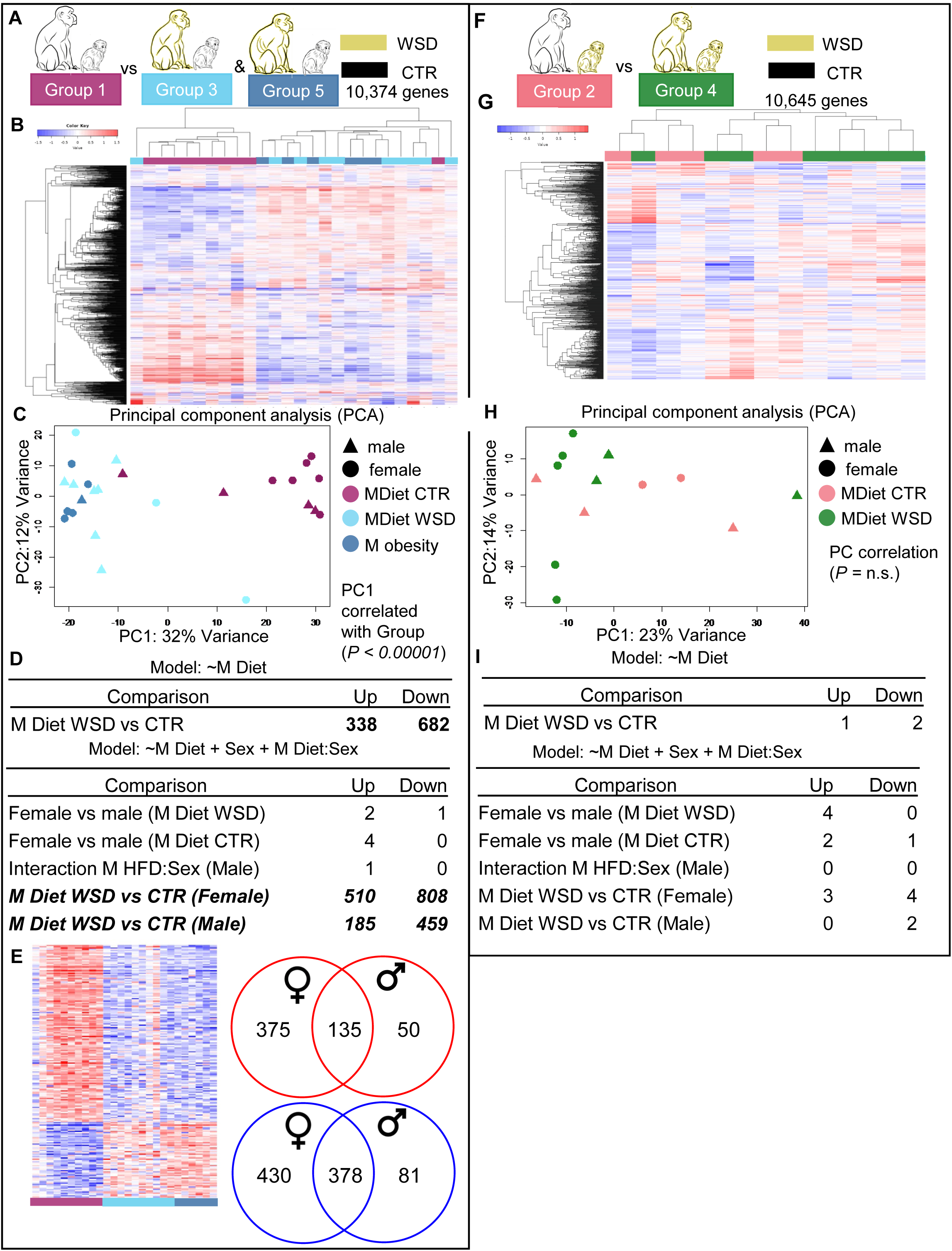
Impact of maternal diet on offspring gene expression with obese and lean dams combined. Colors indicate experimental group with triangles representing males and circles representing females. Panels A-E represent the Group 1 vs Group 3 &5 comparison. Panels F-I represent the Group 2 vs Group 4 comparison. **(A and F)** Group composition and the number of genes retained post-filtering. **(B and G)** Hierarchical clustering of the top 1,000 differentially expressed genes. **(C and H)** PCA of filtered transcriptional profiles. **(D and I)** Linear models and number of significantly differentially expressed genes by comparison. **(E)** Heat map of significantly differentially expressed genes based on maternal diet and venn-diagram showing overlap between significantly up- or downregulated genes across the sexes. Consistent with the heat maps red indicated upregulation and blue indicated down regulation.

We also investigated impacts of M-diet by sex with our increase in sample size by inclusion of Group 5. We fit the data to a linear model accounting for M-diet, sex, and an interaction effect between M-diet and sex. We did not identify strong signals of differential expression between sexes (**Figure 3D**) and instead only identified a small number of genes differentially expressed based on sex alone. Interestingly, we identified strong patterns of differential expression in both male and female offspring in CTR/CTR vs. WSD/CTR (Group 1 vs Group 3 & 5). We identified 510 significantly upregulated and 808 significantly downregulated genes when comparing M-WSD vs. M-CTR diet. In males fed a PW-CTR diet, we identified 185 significantly upregulated and 459 significantly downregulated genes when comparing M-WSD vs. M-CTR diet (**Table S4**). We did not detect an interaction effect likely due to limited sample sizes (**Figure 3D**).

We next tested if males and females exhibited the same changes in expression by M-diet by looking at overlapping sets of differentially expressed genes between the sexes (Group 1 vs Group 3&5). First, significantly differentially expressed genes in offspring fed PW-CTR diets were visualized in a heat map for both sexes (**Figure 3E**). We did not identify any genes significantly upregulated in one sex that were significantly downregulated in the other, but instead detected significant differential expression in many genes only in one sex (data not shown).

We then compared PW-WSD offspring groups exposed to M-CTR or M-WSD (Group 2 vs. Group 4) (**Figure 3F**). Unlike data from PW-CTR offspring, we did not, observe distinct grouping when comparing CTR/WSD vs. WSD/WSD (i.e., Group 2 vs. Group 4) (**Figure 3G**). Additionally, in the Group 2 vs. Group 4 comparison, there was no correlation with any PC (P > 0.05) (**Figure 3H**). When fitting these data to a linear model accounting for M-diet, we identified only 1 significantly upregulated and 2 significantly downregulated genes in the PW-WSD groups (Group 2 vs. Group 4) (**Figure 3I; Table S5**).

### Post-weaning WSD feeding significantly impacts gene expression

To assess the impact of PW-diet on skeletal muscle from male and female offspring respectively, we first compared offspring fed a PW-WSD to those on a PW-CTR within the M-CTR Groups (Group 1 vs. Group 2) (**Figure 4A**). We again performed hierarchical clustering of the top 1,000 differentially expressed genes – regardless of significance – in each group pair to assess partitioning of individuals. We observed clear separation in clustering of individuals from Group 1 vs. Group 2, indicating a potential impact of PW diet on individuals from M-CTR dams (**Figure 4B**). PCA mirrored these findings with PC1 significantly correlated with PW-diet group in the Group 1 vs. Group 2 comparison (*P* = 0.012) accounting for 33% of the observed variance (**Figure 4C**). These findings were further supported by fitting our data to a linear model accounting for PW-diet. When comparing PW-diet treatments in individuals from dams fed a M-CTR (Group 1 vs. Group 2) diet, we identified 287 significantly upregulated and 497 significantly downregulated genes (**Figure 4D**). Unfortunately, limited sample sizes prevented testing for of differential expression by sex as we only had two females in the CTR/WSD group. Upon further investigation, however, we found that both females and a single male in the CTR/WSD group (Group 2) exhibited elevated fasting insulin levels (**Table S1**) consistent with previous studies^16^ and that those with the highest fasting insulin (**Table S1**) exhibited the most extreme differential expression (**Figure 4E**).

**Figure 4.**
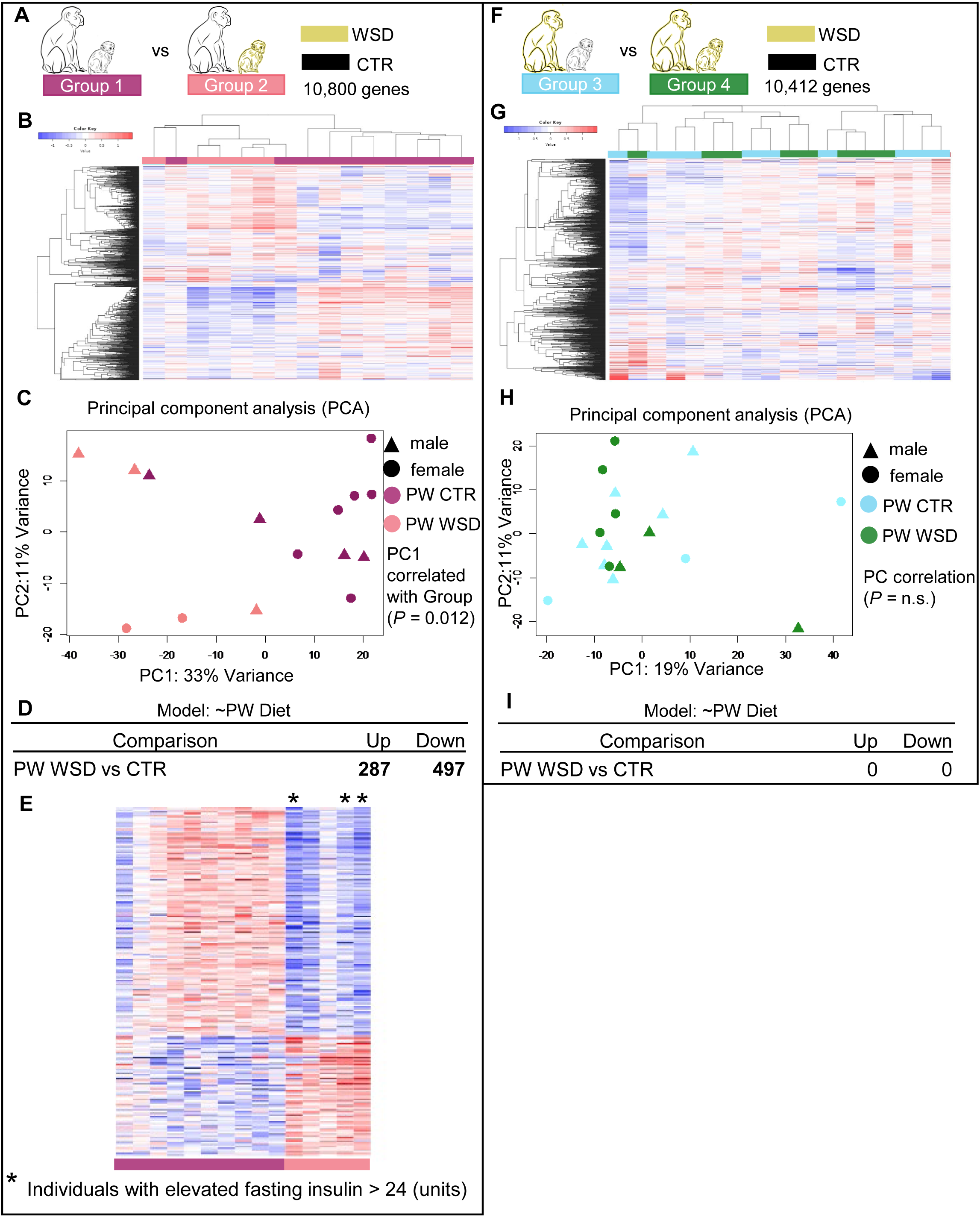
Impact of post-weaning diet on offspring gene expression. Colors indicate experimental group with triangles representing males and circles representing females. Panels A-E represent the comparison of Group 1 vs Group 2. Panels F-I represent the comparison of Group 3 vs Group 4. **(A and F)** Group composition and the number of genes retained post-filtering. **(B and G)** Hierarchical clustering of the top 1,000 differentially expressed genes. **(C and H)** PCA of filtered transcriptional profiles. **(D and I)** Linear models and number of significantly differentially expressed genes by comparison. **(E)** Heat map of significantly differentially expressed genes based. Asterisks indicate individuals with elevated fasting insulin.

We also compared offspring fed a PW-WSD to those on a PW-CTR within the M-WSD groups (Group 3 vs. Group 4) (**Figure 4F**). We did not observe distinct clusters comparing Group 3 vs. Group 4, no PC significantly correlating with group (P > 0.05) (**Figure 4G**) and no differentially expressed genes were identified (**Figure 4I**) suggesting PW-diet had little or no effect on offspring gene expression profiles when offspring were exposed to M-WSD.

### Female offspring exhibit interaction effects between maternal and post-weaning diets

To test for interaction effects between PW and M diets, we combined all data from offspring born to lean mothers (Groups 1-4) (**Figure 5A**). Using PCA, we identified distinct clustering by Group with PC1 significantly correlated with experimental group (*P* = 0.00007) accounting for 28% of the variance (**Figure 5B**). When we fit these data to a linear model accounting for M-diet, PW-diet, and interaction between PW and M diets, we identified 121 significantly upregulated and 213 significantly downregulated genes due to the interaction between diets (**Figure 5C**). When visualized as a heat map, there were clear cases of individuals not fitting the pattern of DE between Group 1 (CTR/CTR) and the other three groups (CTR/WSD; WSD/CTR; WSD/WSD) (**Figure 5D**). Combined, previous findings of increased variance in males by M-diet, correlation of read count with group in this comparison, and the presence of outliers in our DE heatmap led us to assess impacts of diet interactions by sex.

**Figure 5.**
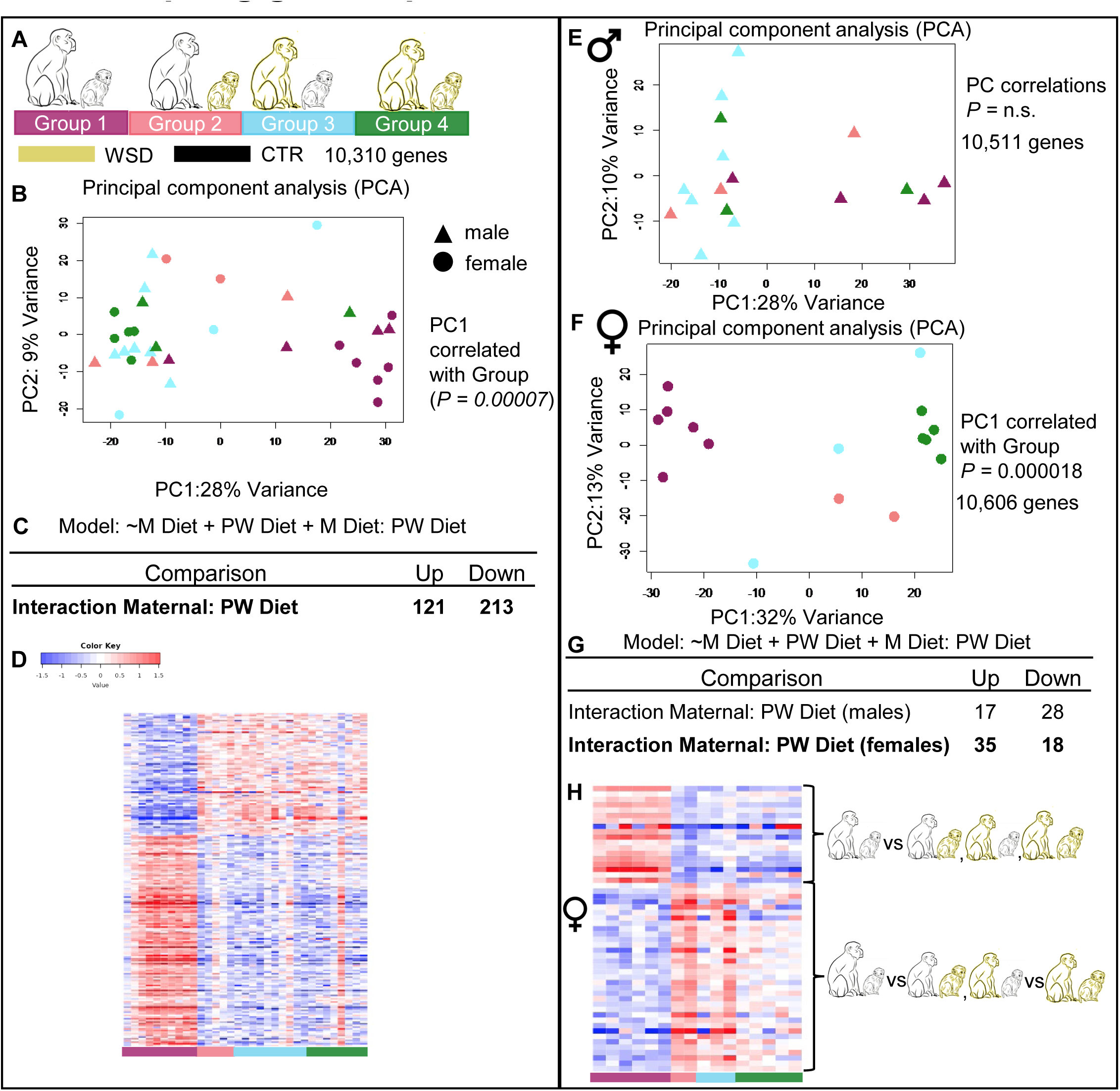
Interaction effects between maternal and post-weaning diets on offspring gene expression. Colors indicate experimental group with triangles representing males and circles representing females. **(A)** Group composition and the number of genes retained post-filtering. **(B)** PCA of filtered transcriptional profiles with sexes combined. **(C)** Linear model and number of significantly differentially expressed genes. **(D)** Heat map of significantly differentially expressed genes with sexes combined. **(E)** PCA of male filtered transcriptional profiles. **(F)** PCA of female filtered transcriptional profiles. **(G)** Linear model and number of significantly differentially expressed genes in males and females separately. **(H)** Heat map of significantly differentially expressed genes in female offspring. Brackets indicate groupings of observed patterns of differential expression (CTR/CTR vs CTR/WSD or WSD/CTR or WSD/WSD) and (CTR/CTR vs CTR/WSD or WSD/CTR vs WSD/WSD).

When separating groups by sex, we identified striking patterns of dissimilarity. In males, no clustering of groups or significant correlations with any PC were observed (*P* > 0.05) (**Figure 5E**). In contrast, females exhibited clear clustering by group and a significant correlation of Group with PC1 (*P = 0.*00018) (**Figure 5F**). We then fit a linear model accounting for M-diet, PW-diet, and an interaction effect between M and PW- diets (**Figure 4G**). While we did not identify many significantly differentially expressed genes (35 significantly upregulated and 18 significantly downregulated in females), we did observe a clear clustering pattern in the heatmap of differentially expressed genes (**Figure 5H**). In the heatmap, gene clusters presented in three groups (CTR/CTR vs CTR/WSD or WSD/CTR or WSD/WSD) instead of the four treatment groups present in the experimental design (**Figure 5D**). In female offspring, there were two different patterns of differential expression. In the 18 significantly downregulated genes, patterns were consistent with those observed with sexes combined; exposure to WSD either in M-diet or PW-diet resulted in downregulation compared to the CTR/CTR group. In the 35 significantly upregulated genes, however, individuals exposed to WSD *either* in PW or M diet – but not both – exhibited the strongest upregulation. Individuals from the WSD/WSD group exhibited patterns of expression that were more similar to the CTR/CTR group (**Figure 5H**).

### Top gene analysis and KEGG enrichment of differentially expressed genes reveals targets of interest

Differential expression analyses based on M-diet in offspring fed PW-CTR yielded the strongest signal (Group 1 vs Groups 3 & 5). To biologically assess the impacted genes/gene groups, we identified the most differentially expressed genes in all animals and in each sex using (**Table 1**). While individual gene dysregulation can have major biological consequences, we also performed KEGG pathway enrichment to identify biological pathways enriched for differentially expressed genes. We used raw log-fold changes to identify pathways significantly up- or downregulated. We performed this analysis on sexes separate and sexes combined (**Figure 6**). Top KEGG pathways included those pathways predicted to be different based on our prior work like insulin signaling, PI3K-AKT signaling, metabolic pathways and fatty acid degradation. Female offspring had a stronger influence on enrichment in these pathways than male offspring.

**Figure 6.**
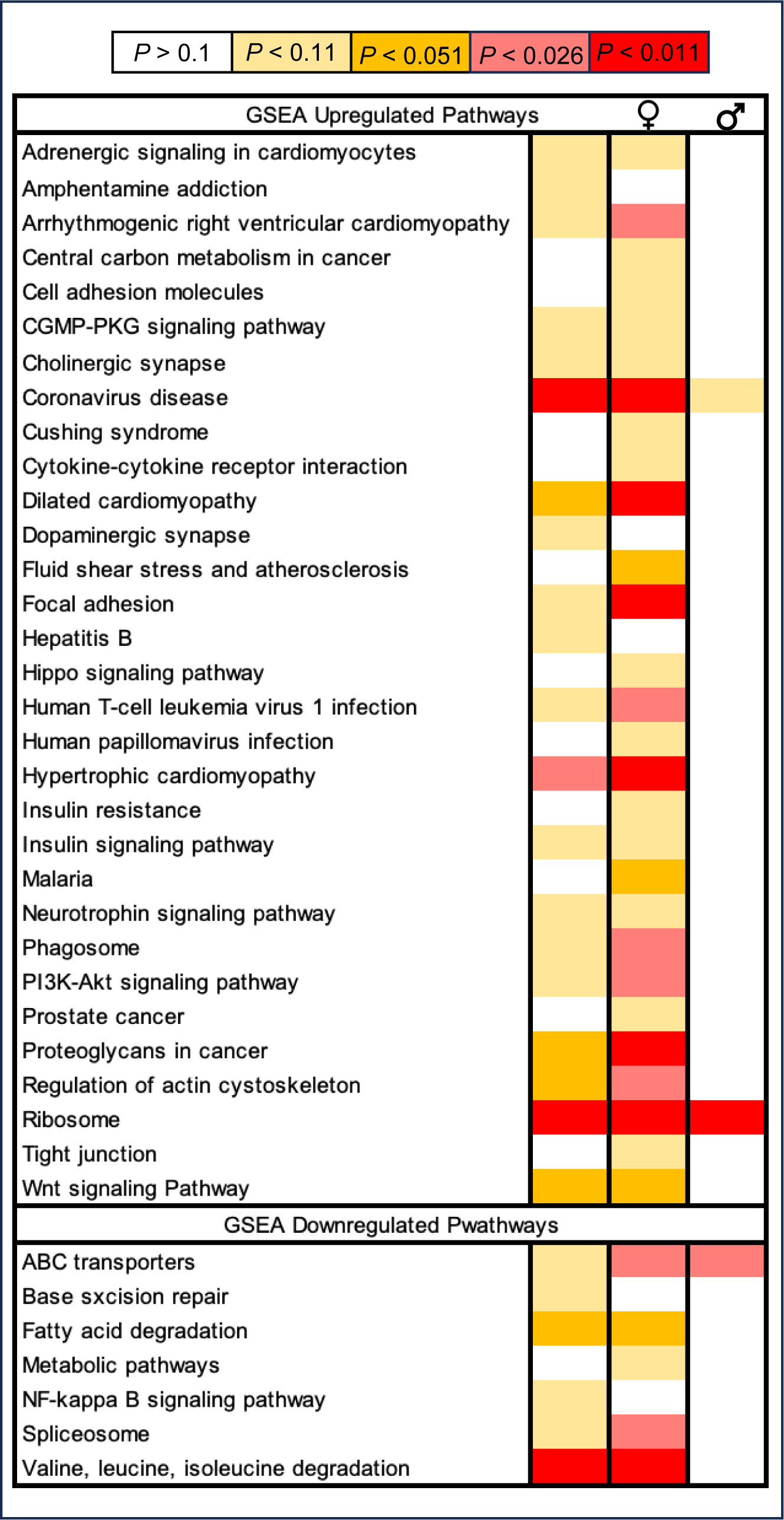
Heatmap of KEGG Pathway enrichment based on maternal diet. Heat map represents KEGG pathways enrichment for significantly differentially expressed genes. GSEA methods used for identification of upregulated and downregulated pathways. KEGG pathways are represented alphabetically. Heat map columns represent sexes combined, females only, and males only. Intensity correlates with significance level.

**Table 1.** Top 15 Differentially Expressed Genes by Maternal Diet. Comparison of Group 1 vs Groups 3 & 5 combined. Subsections separated by sexes combines, females only, and males only. Only elements with gene names included

| Sexes Combined |  |  |  |  |  |
| --- | --- | --- | --- | --- | --- |
| Gene Name | log2fold Δ | Adj P-Value |  |  |  |
| *ifi27* | -3.17 | 3.06E-33 |  |  |  |
| *gadd45g* | -2.31 | 9.16E-16 |  |  |  |
| *itih4* | -2.14 | 3.43E-09 |  |  |  |
| *pla2g4b* | -2.13 | 1.79E-14 |  |  |  |
| *leng8* | -2.04 | 7.33E-18 |  |  |  |
| *hsf4* | -1.98 | 6.91E-16 |  |  |  |
| *pan2* | -1.98 | 1.97E-20 |  |  |  |
| *otud1* | 1.97 | 1.38E-03 |  |  |  |
| *map3k6* | -1.97 | 4.96E-19 |  |  |  |
| *neil1* | -1.88 | 6.86E-13 |  |  |  |
| *scarf1* | -1.86 | 3.28E-18 |  |  |  |
| *tmem82* | -1.79 | 1.39E-12 |  |  |  |
| *clasrp* | -1.75 | 2.16E-15 |  |  |  |
| *slc7a6* | -1.74 | 1.70E-18 |  |  |  |
| *znf7* | -1.73 | 2.65E-18 |  |  |  |
| Females Only |  |  | Males Only |  |  |
| Gene Name | log2fold Δ | Adj P-Value | Gene Name | log2fold Δ | Adj P-Value |
| *ifi27* | -3.27 | 4.49E-15 | *ifi27* | -2.99 | 1.10E-08 |
| *myh4* | -3.09 | 1.02E-03 | *spp1* | -2.84 | 4.95E-04 |
| *otud1* | 2.57 | 2.25E-03 | *gadd45g* | -2.01 | 6.39E-05 |
| *gadd45g* | -2.43 | 6.20E-10 | *leng8* | -1.92 | 4.89E-06 |
| *itih4* | -2.28 | 5.71E-06 | *hsf4* | -1.86 | 2.12E-05 |
| *plag4b* | -2.25 | 2.55E-09 | *dsn1* | -1.85 | 1.97E-05 |
| *map3k6* | -2.08 | 1.72E-11 | *plxna3* | -1.85 | 6.59E-04 |
| *pan2* | -2.08 | 1.82E-13 | *pla2g4b* | -1.82 | 1.74E-04 |
| *leng8* | -2.07 | 3.15E-10 | *map3k6* | -1.76 | 8.03E-06 |
| *hsf4* | -2.02 | 4.37E-09 | *itih4* | -1.76 | 8.08E-03 |
| *map3k8* | -2.01 | 3.15E-10 | *pan2* | -1.72 | 2.51E-06 |
| *neil1* | -1.97 | 1.97E-08 | *nphp4* | -1.71 | 5.77E-05 |
| *znf460* | 1.96 | 1.13E-12 | *scarf1* | -1.7 | 3.49E-06 |
| *slc25a1* | -1.91 | 2.98E-08 | *clasrp* | -1.7 | 1.42E-05 |
| *slc7a6* | -1.9 | 2.17E-14 | *mpz* | -1.66 | 5.30E-03 |

## DISCUSSION

Fetal nutrient and growth factor availability during pregnancy significantly impacts muscle size, fiber composition, and programs responsiveness of pathways important for muscle growth, repair, and metabolism^25,26^ in mammals and avian species. Maternal diets that are both calorically dense^26–28^ or calorie/protein insufficient^29–31^ during pregnancy impair fetal myogenesis, resulting in smaller skeletal muscle fibers and reduced downstream insulin signaling years after birth. Our previous findings in a Japanese macaque model found impaired skeletal muscle metabolic flexibility and reduced insulin-stimulated glucose uptake in fetal offspring from dams consuming a WSD^19–21^. Here, we find that persistent shifts in transcriptional response may underlie these programmed changes in offspring skeletal muscle function and increase the risk of metabolic disease.

Our differential expression analyses revealed several interesting and somewhat unexpected findings. First, maternal diet was a major driver of transcriptional changes not maternal obesity (**Figure 2**; **Figure 3**). Given multigenerational studies of maternal obesity in rodent models^32^, we anticipated finding transcriptional changes but hypothesized that maternal obesity and insulin resistance, more so than maternal WSD, would significantly impact offspring transcriptome in skeletal muscle. Evidence from several epidemiological studies and clinical cohorts strongly supports that maternal obesity, potentially mediated through poor metabolic health status^33–35^, increases infant adiposity and risk of offspring obesity^1,36,37^. Additionally, in a cohort of greater than 30,000 adult offspring, maternal obesity was linked to higher all-cause mortality and greater risk of cardiovascular disease in adult offspring^38^. Therefore, the persistent impact of maternal WSD in the absence of maternal obesity or maternal insulin resistance on the skeletal muscle gene expression, particularly when offspring were switched to a healthy control diet, was unexpected (**Figure 2**). More so, the lack of a further effect of maternal obesity as compared to maternal WSD alone on offspring gene expression suggests a poor quality WSD is sufficient for metabolic reprogramming in offspring skeletal muscle, at least in a nonhuman primate model (**Figure 3**). There are two caveats to these findings. First, given our small sample size, the lack of difference with maternal obesity may be due to Type II error. Further exploration with an expanded sample sizes is needed to thoroughly address the potential impacts of maternal body composition on offspring health. Additionally, sex distribution was not matched in this analysis. Given other analyses showed difference in response between males and females, the unbalanced sex distribution between the two groups may obscure the findings.

Transcriptomic studies in liver, skeletal muscle and adipose tissue of fetal; or adult offspring in mouse models identify differing transcriptional responses of male and female offspring to maternal obesity^39^, offering a probable explanation for sexually dimorphic metabolic outcomes as male offspring have been reported to have worse metabolic adaptations than their female counterparts^39–43^; however, other studies show worse outcomes in female offspring^44,45^. While these studies in rodents suggest fetal exposure to maternal obesity alters offspring metabolism through transcriptional mechanisms, factors such as degree of high fat diet exposure, housing temperature, developmental stage/maturity at birth, and litter births may limit the relevance to human populations. In our study, we also observed that male and female offspring were impacted differently by M-diet, with females exhibiting more differentially expressed genes than males, but we were surprised to see few genes differentially expressed based on sex alone (**Figure 3**). Increased transcriptional responses in females suggests fetal adaptations to M-WSD allows for genetic and molecular compensations that may support greater resiliency to later life environmental stressors compared to male offspring. In male offspring, we observed increased variance that was most clearly demonstrated when assessing interaction effects between M and PW-diets. Again, these inferences are not without caveats. Due to the limited sample size, we likely did not have the statistical power to detect an interaction effect between treatment and sex. To understand the differences in males and females, we relied on general global patterns of expression differing between the sexes, and differences in significance across PC space to ascertain different responses between the sexes. Additionally, given our observation that male and female offspring are affected differently by M-diet, it is surprising that we have not yet observed differences in functional outcomes measures including insulin-stimulated glucose uptake^19^, mitochondrial oxidative metabolism^20,21^, in offspring skeletal muscle but speculate that dysregulation of different pathways may lead to the same outcome. Thus, it is important for future work to account for sex when assessing mechanisms for how maternal health or diet impacts on offspring.

In metabolically healthy individuals, switching to a higher fat WSD leads to adaptations in skeletal muscle to promote fatty acids uptake, storage and oxidation in skeletal muscle^46,47^, in part, through transcriptional coordination of PPARs, PGC-1α and ERα signaling^48–50^. In contrast, those with metabolic disorders, such as obesity, insulin resistance, and type 2 diabetes, demonstrate reduced mitochondrial content and diminished metabolic flexibility^51,52^ potentially driven by a blunted transcriptional response^52^. In our study, skeletal muscle from offspring of M-CTR fed dams had a transcriptional shift in gene expression, but there was little to no transcriptional response based on PW-diet in offspring of M-WSD dams (**Figure 4**). The lack of a robust response in lean, healthy adolescent offspring from M-WSD dams is unexpected indicating that M-WSD suppresses later transcriptional response to WSD. This may be due to either an early life activation and programming of these pathways such that no additional adaptation is required, or perhaps M-WSD leads to a down-regulation of nutrient sensing signals like AMPK, SIRT1 and mTOR^53^ resulting in suppressed metabolic fuel shifts as noted in adults with obesity. Regardless, these results suggest that returning to a healthy chow diet (a postweaning diet intervention) may be insufficient in mitigating the effects of poor maternal diet.

An additional trend worth highlighting is the stark differences in interaction effects between M and PW- diets observed in male vs. female offspring respectively (**Figure 5**). When assessing interaction effects directly, we see clear clustering of individuals based on treatment group in female but not male offspring. In our analysis, female offspring in the WSD/WSD group showed a distinct pattern of gene response that was different from either the M-WSD or PW-WSD group alone. This suggests M-WSD may create a transcriptional pattern that “primes” muscle to be compatible with continued WSD exposure and may accelerate disease risk. In contrast, a mismatch between maternal and postweaning diet leads to unique transcriptional pattern from that observed with the matched maternal and postweaning diet. Further investigation of these distinct gene sets may reveal mechanisms or biomarkers for individuals with increased risk.

While we were most interested in general patterns observed in this study – namely the impact of treatments (i.e. M-obesity, M-diet, and PW -diet) on transcription globally, there are a few interesting genes and gene pathways identified that warrant further investigation in future studies. As M-diet exhibited the most major effect on offspring gene expression, we investigated the top differentially expressed, and KEGG enriched pathways significantly impacted by this treatment in both sexes independently and combined (**Table 1**; **Figure 6**). Within the group of top 15 differentially expressed genes, were several involved in stress-responsive mitogen-activated protein kinases (MAPK) signaling pathway, *map3k6*, *map3k8*, and *pla2g4b*. In skeletal muscle, activation of MAPK, most notably p38 MAPK, and c-Jun NH2-terminal kinase (JNK)^54^ but also TPL2 (i.e. *map3k8)*^55^ is linked to insulin resistance. As noted, we have observed impaired glucose uptake in offspring skeletal muscle, which was not linked to changes in insulin action typically associated with muscle insulin resistance observed in obesity. Changes in MAPK may be a novel mechanism to explain offspring insulin resistant phenotype. Downregulation of *ifi27* (IFN-α inducible protein 27) and *gadd45g* (growth arrest and DNA damage inducible gamma) was most also consistent across sexes based on M-diet, with *ifi27* exhibiting the most extreme downregulation in both sexes. In adipocytes, IFI27 localizes to inner mitochondria matrix, and promotes fatty acid oxidation through interactions with the trifunctional protein (HADHA)^56,57^. This is especially interesting as we have observed impaired fat oxidation in skeletal muscle from M-WSD offspring^21^.The downregulation of *gadd45g* was particularly surprising as it is typically upregulated due to stress^58^, and in fetal muscle, we observed *gadd45g* was upregulated in response to maternal obesity^20^. The opposite augmentation in expression due to maternal WSD in the postweaning period may be an adaptation that leaves muscle cells more vulnerable to oxidative damage and may promote a proinflammatory phenotype as demonstrated in hematopoietic stem and immune progenitor cells in these offspring^13^.

At the genetic pathway level, there were also some interesting trends (**Figure 6**). Differentially expressed genes were enriched in KEGG pathways for “Ribosome” in both sexes independently and combined resulting in significant upregulation of ribosomes. We also noted enrichment in downregulation of “Valine, leucine, isoleucine degradation”. This is not surprising as skeletal muscle is in essence the primary storage site for amino acids. This also aligns with dysregulation of mTOR signaling which is commonly observed in aged- or insulin resistant muscles^59,60^. Upregulation of “Ribosomes” combined with downregulation of “Valine, leucine, isoleucine degradation” and with downregulation of “Spliceosomes” – also observed – suggests a potential breakdown of transcription-translation feedback loops which could be impacting expression of other gene pathways. We also noted upregulation of several pathway involved in glucose uptake including “PI3K-Akt signaling”, “insulin resistance”, and “insulin signaling” with M-WSD as well as upregulation of “CGMP-PKG signaling pathway” which is responsive to NO and is thought to mediates glucose uptake in basal or contraction-stimulated conditions^61,62^. The upregulation of these pathways may be compensatory to the reduced insulin responsiveness measured in fetal and juvenile skeletal muscle of offspring exposed to M-WSD and/or may be elevated due to the potential breakdown in transcription-translation feedback loops. Alternatively, there may be an upregulation in genes within these pathways that negatively regulate insulin action. Overall, these data indicate a persistent impact of maternal diet on offspring skeletal muscle transcriptome and regulation of glucose metabolism.

In summary, our results show that a maternal Western-style diet, even in the absence of maternal obesity and insulin resistance, is sufficient to reprogram offspring skeletal muscle transcriptional response in male and female offspring in a Japanese macaque model. The effects on gene expression were robust lasting years after exposure and influenced offspring responsiveness to a postweaning western-style diet that was sex-dependent. We propose that this persistence in transcriptional response based on maternal diet contributes to the observed increased susceptibility to metabolic diseases in offspring. Future work will be aimed at identifying the cellular mechanisms leading to persistent changes in gene networks.

## METHODS

### Animals

All animal procedures were approved by and conducted in accordance with the Institutional Animal Care and Use Committee of the Oregon National Primate Research Center (ONPRC) and Oregon Health and Science University. The ONPRC abides by the Animal Welfare Act and Regulations enforced by the USDA and the Public Health Service Policy on Humane Care and Use of Laboratory Animals in accordance with the Guide for the Care and Use of Laboratory Animals published by the NIH.

### Experimental Model

Adult female Japanese macaques were group housed in indoor/outdoor pens with 1-2 males and were fed ad libitum either a CTR diet fed (15% calories from fat originating primarily from soybeans and corn; Monkey Diet no. 5000; Purina Mills) or WSD (37% calories from fat primarily from corn oil, egg, and animal fat; TAD Primate Diet no. 5LOP, Test Diet, Purina Mills) for at least 1.5 years prior to pregnancy. Carbohydrate content differed between the two diets, with sugars (mainly sucrose and fructose) constituting 19% of the western-style diet but only 3% control diet. Monkeys on the WSD were also given calorically dense treats (35.7% of calories from fat, 56.2% of calories from carbohydrates, and 8.1% of calories from protein) once daily. A subset of dams in the WSD group remained lean and insulin-sensitive similar to females fed a CTR diet while the others became obese (**Table S2**). Adult females were classified as lean or obese based on percentage body fat obtained by dual-energy X-ray absorptiometry (DEXA) in the fall preceding the pregnancy of interest^20^. Adult females in the WSD group with increased body fat (M-obesity) were older, had been fed the WSD for longer and were more insulin resistant (**Table S2**). An intravenous glucose tolerance test used to assess insulin sensitivity prior to pregnancy determination (fall) and during the 3^rd^ trimester as previously described^63^. Only singleton births were included in the cohort.

Offspring were born naturally and remained in their home colony until weaning. At 7-8 months of age, juvenile offspring were weaned to new group housing with 6–10 similarly aged juveniles from both maternal diet groups and 1–2 adult females. These new housing groups were then fed either CTR or WSD, creating four offspring groups. Body weight and body composition by DEXA was taken at ∼38 months of age. Insulin sensitivity was measured by IV GTT at 36 months as previously described^16^. Sample size for each group (maternal diet/offspring diet) included 12 CTR/CTR (7 female [F], 5 male [M]), 5 CTR/WSD (2 F, 3 M), 13 WSD/CTR (5 F, 8 M), and 8 WSD/WSD (5 F, 3 M), juvenile offspring from 9 CTR dams and 14 WSD dams were included in this study. In male or female offspring, there was no increase in body weight or percent body fat across groups with PW-WSD(**Table 2**); however, as previously shown in larger cohorts of offspring^16^, male and female CTR/WSD offspring had higher fasting insulin and higher insulin area under the curve during iv GTT as compared to CTR/CTR(**Table 2**). In females, fasting insulin and insulin AUC was also higher in CTR/WSD compared to WSD/WSD (**Table 2**).

**Table 2.** Male and female offspring physiology at 3 years of age.

|  |  | CTR/CTR<br>(Group 1) | CTR/WSD<br>(Group 2) | WSD/CTR<br>(Group 3) | WSD/WSD<br>(Group 4) | Obese/CTR<br>(Group 5) |  |  |  |
| --- | --- | --- | --- | --- | --- | --- | --- | --- | --- |
| Offspring<br>number | Female | 6 | 2 | 3 | 5 | 5 | Main effects |  |  |
|  | Male | 4 | 3 | 7 | 3 | 2 | M | PW | INT |
| Weight<br>(g) | All | 5937 ± 263 | 5813 ± 360 | 6177 ± 302 | 5910 ± 275 | 6047 ± 99 | ns | ns | ns |
|  | F | 5748 ± 423 | 5531 ± 298 | 5676 ± 729 | 5628 ± 341 | 6063 ± 114 | ns | ns | ns |
|  | M | 6220 ± 162 | 6002 ± 599 | 6391 ± 306 | 6379 ± 378 | 6005 ± 270 | ns | ns | ns |
| Lean<br>Mass (g) | All | 4772 ± 217 | 4745 ± 271 | 4985 ± 249 | 4880 ± 221 | 4836 ± 127 | ns | ns | ns |
|  | F | 4607 ± 348 | 4469 ± 285 | 4515 ± 588 | 4681 ± 244 | 4915 ± 114 | ns | ns | ns |
|  | M | 5018 ± 122 | 4928 ± 418 | 5187 ± 256 | 5213 ± 407 | 4640 ± 395 | ns | ns | ns |
| Body Fat<br>(%) | All | 15.6 ± 0.7 | 14.1 ± 1.0 | 15.1 ± 0.5 | 13.2 ± 1.0 | 15.9 ± 1.1 | ns | .04 | ns |
|  | F | 15.8 ± 1.2 | 15.0 ± 0.9 | 15.9 ± 1.1 | 12.5 ± 1.0 | 14.8 ± 0.8 | ns | ns | ns |
|  | M | 15.3 ± 0.5 | 13.5 ± 1.6 | 14.7 ± 0.6 | 14.3 ± 2.0 | 18.5 ± 3.1 | ns | ns | ns |
| Insulin<br>(μU/mL) | All | 6.7 ± 1.6<sup>a</sup> | 20.9 ± 6.0<sup>d</sup> | 5.3 ± 1.0 | 9.9 ± 1.1 | 5.7 ± 1.3 | .009 | .0002 | .04 |
|  | F | 7.9 ± 2.6<sup>a</sup> | 28.4 ± 10.2<sup>d</sup> | 7.8 ± 2.9 | 9.3 ± 1.8 | 6.7 ± 1.6 | .02 | .02 | .01 |
|  | M | 5.0 ± 0.4<sup>a</sup> | 16.0 ± 7.4 | 4.2 ± 0.5 | 10.8 ± 0.7 | 3.2 ± 1.0 | ns | .006 | ns |
| Glucose<br>(mg/dL) | All | 55 ± 5 | 60 ± 4 | 55 ± 3 | 55 ± 1 | 48 ± 2 | ns | ns | ns |
|  | F | 50 ± 6 | 63 ± 10 | 51 ± 5 | 55 ± 2 | 48 ± 3 | ns | ns | ns |
|  | M | 62 ± 6 | 59 ± 4 | 56 ± 4 | 56 ± 2 | 47 ± 4 | ns | ns | ns |
| Insulin<br>AUC | All | 1578 ± 213 | 2364 ± 345 | 1738 ± 222 | 2087 ± 190 | 1797 ± 398 | ns | .03 | ns |
|  | F | 2036 ± 364<sup>a</sup> | 3161 ± 192<sup>d</sup> | 2135 ± 288 | 1868 ± 135 | 1410 ± 169 | ns | ns | .02 |
|  | M | 1256 ± 109 | 1833 ± 180 | 1567 ± 279 | 2451 ± 410 | 2765 ± 1316 | ns | .03 | ns |
| Glucose<br>AUC | All | 10095 ± 467<sup>c</sup> | 8493 ± 449 | 8226 ± 444 | 8263 ± 427 | 8440 ± 561 | .04 | ns | ns |
|  | F | 9915 ± 582 | 7637 ± 815 | 8392 ± 372 | 7923 ± 558 | 8587 ± 779 | ns | ns | ns |
|  | M | 10320 ± 843 | 9064 ± 207 | 8154 ± 631 | 8830 ± 638 | 8075 ± 601 | ns | ns | ns |

Data is shown as the mean ± SEM. For each measurement, group data is given for all animals, and then separated by sex. Fasting glucose and insulin were measured in anesthetized animals. Body composition was measured by DEXA and percent fat calculated from total mass. Data were analyzed by two-way ANOVA for main effects of maternal (M) or postweaning (PW) diet or for an Interaction (INT) across groups 1-4. When significant (p<0.05), post hoc comparison between groups is indicated by letters: <sup>a</sup>group 1 vs. 2; <sup>b</sup>group 3 vs. 4; <sup>c</sup>group 1 vs. 3; <sup>d</sup>group 2 vs. 4. A Student's t- test was used to compare offspring in group 3 and group 5; \* p<0.05. ns, not significant.

An initial concern with our experimental design was that variables unrelated to experimental group, and their main variables, could be influencing transcriptional profiles, specifically, the cohort year in which offspring were reared and the genetic relatedness of individuals (**Table S1**). For each cohort year (**Table S1**), we observed a complete span of PC1 space, with no significant correlations between any PC and cohort year (**Figure S4**).

Importantly, while there is some variability in relatedness of our combined 40 Japanese macaque offspring, all individuals are closely related. In all but one case, dams reared one or two offspring with one dam rearing three offspring (**Table S1**). Offspring were randomized across treatment groups (i.e., one dam may have reared two offspring in the same treatment group or in different treatment groups). The identity of sires is unknown; however, the family structure of Japanese macaques limits the number of possible sires so in all cases there are one or two sires possible within each harem. This means all offspring are likely at least half-siblings.

### Juvenile muscle collection

Juvenile animals were sacrificed between 37 and 40 months of age with an average age of 38 months for all groups. At time of necropsy, juvenile skeletal muscles including gastrocnemius, soleus, vastus lateralis, and rectus femoris were rapidly dissected of fascia and portions were flash frozen in liquid nitrogen. Frozen muscle was pulverized was pulverized on dry ice and stored at –80°C until analysis.

### RNA Extraction

RNA was extracted from approximately 50 mg of pulverized gastrocnemius using PureLink RNA mini kit (Thermofisher, Waltham, MA, USA) according to manufacturer’s instructions except homogenization in trizol was performed using an Omni bead mill with a CryoCool system. Each cryomill tube had 5 x 2.8mm ceramic beads (Omni Inc, Kennesaw, GA, USA) and was run for 2 x 30 secs at 6 m/s with a 10 sec dwell period.

### RNA-seq Data Processing and Normalization

RNA-seq library preps were generated using the Kapa stranded mRNA-seq kit (Roche, Basel, Switzerland). Samples were individually barcoded sequenced on the Illumina Hi-Seq 4000 (Illumina Inc, San Diego, CA, USA). Sequencing reads were demultiplexed via Illumina’s bcl2fastq software (Illumina Inc, San Diego, CS, USA) and trimmed and quality filtered using BBDuk (BBMap-Bushnell B. - sourceforge.net/projects/bbmap/). Reads were aligned to the rhesus macaque reference genome version 8.0.1 using the STAR aligner^64^ version 2.5.4b and read count tables generated using the -quantMode gene counts function in STAR. Preprocessing of read counts was performed with integrated Differential Expression and Pathway analysis (iDEP) version 0.93^65^. This package implements R programming language version 4.05^66^ and Bioconductor^67,68^ version 3.12. Total read counts per gene were input and metadata for each hypothesis tested separately into iDEP93. A cut-off of 0.5 CPM (copies per million) was used to filter genes with low counts for all libraries prior to differential expression analyses. We normalized and log2 transformed gene counts using EdgeR^69^ in IDEP93^65^. When assessing all 40 individuals for read count distributions, we identified a correlation between read count and M-diet (*P =* 0.00657) (**Figure S2**). This correlation did not persist in any of our hypothesis tests and no significant correlations between read count and other experimental variables were found when assessing impacts of PW-diet, M-diet, or M-obesity, (with sexes combined or split). There were also no correlations identified when testing for interaction effects between PW-diet and M-diet when groups were split by sex. However, we did see a significant correlation of read count and Group (*P =* 0.00729) and read count and M-diet (*P* = 0.0067) in testing for PW-diet and M-diet interaction effects with sexes combined.

### Differential Expression Analysis and Pathway Enrichment

iDEP93^65^ was used for differential expression analyses. Multivariate patterns of gene expression between experimental groups were characterized with Principal Component Analysis (PCA). Coordinates were exported from iDEP93 and visualized in RStudio using Base R^66^. Heatmaps were generated using data visualization in IDEP93 implementing hierarchical clustering of the top 1,000 differentially expressed genes. Differential expression analyses were performed with DESeq2^70^ with a minimum fold change of 1.5 and a False Discovery Rate (FDR) of 0.05. We selected main effects for each comparison with interaction effects included when appropriate. We performed three iterations of KEGG Pathway enrichment using Generally Applicable Gene set Enrichment (GAGE) and Gene Set Enrichment Analysis (GSEA) with an FDR of 0.05. KEGG^71^ enrichment was performed using Ensembl Release 100.

## AUTHOR CONTRIBUTIONS

C.E.M. conceptualized and designed the study in collaboration with the NHP consortium including S.R.W., K.M.A., J.E.F., P.K., and M.G. Tissue acquisition was performed at Oregon Health Sciences University led by P.K with T.A.D. Molecular work was performed by B.H. Bioinformatic analyses were performed by E.A.B. with L.N. and D.W.T. Statistical analysis and data visualization were performed by E.A.B. Clayton M. Small provided additional insights on statistical analyses. The manuscript was written by E.A.B and C.E.M. with all authors contributing to the editing of the manuscript.

## DATA AVAILABILITY

All sequencing reads are available on the Sequencing Read Archive (SRA) under Submission ID SUB13474614.

