## supplemental table and figures for "Maternal Western-style diet programs skeletal muscle gene expression in lean adolescent Japanese macaque offspring"

**Supplemental Figure Legends**

**Figure S1. Outlier Identification via Principal Component Analysis.**

PCA of all 40 offspring originally included in the study. Color indicates group association. Triangles indicate males and circles indicate females. Outlier is identified by an asterisk.

**Figure S2. Read count distribution per sample post-filtering.**

Histogram of total read count (millions) per sample. Bars are colored by group/sex affiliation.

**Figure S3. Impact of maternal diet on offspring gene expression.**

Colors indicate experimental group with triangles representing males and circles representing females. **(A)** Group composition and the number of genes retained post-filtering. **(B)** Hierarchical clustering of the top 1,000 differentially expressed genes. **(C)** PCA of filtered transcriptional profiles. (**D)** Linear models and number of significantly differentially expressed genes by maternal diet.

**Figure S4. Principal component analysis based on cohort year.**

Colors indicate year individuals were born.

### Figure S1. Outlier identification via Principal Component Analysis.

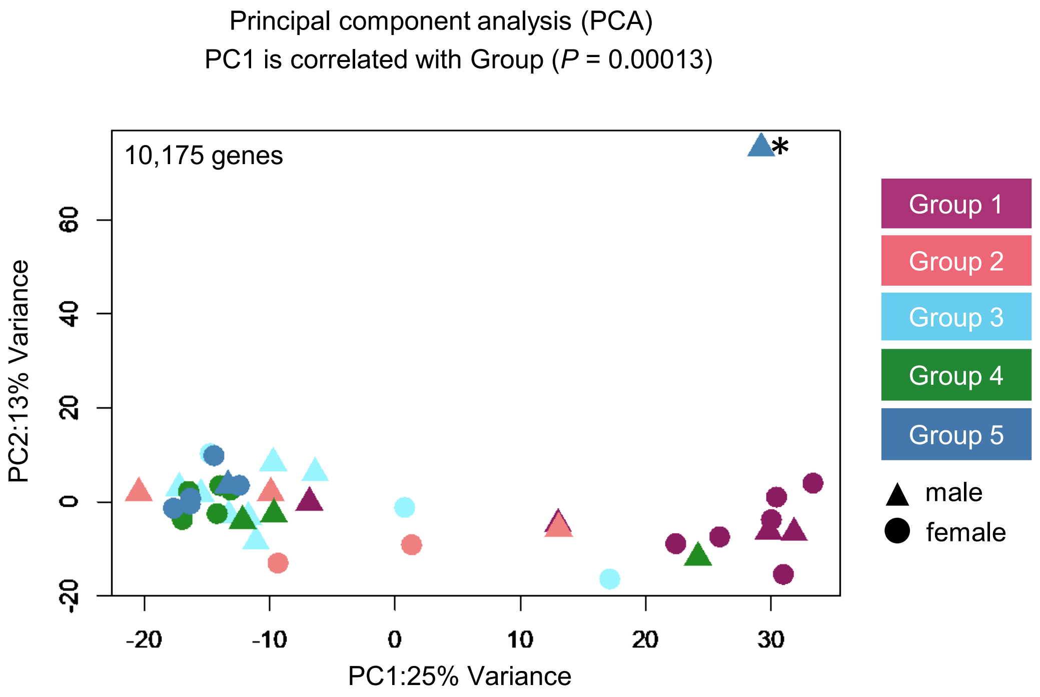

#### Figure S2. Read count distribution per sample post-filtering.

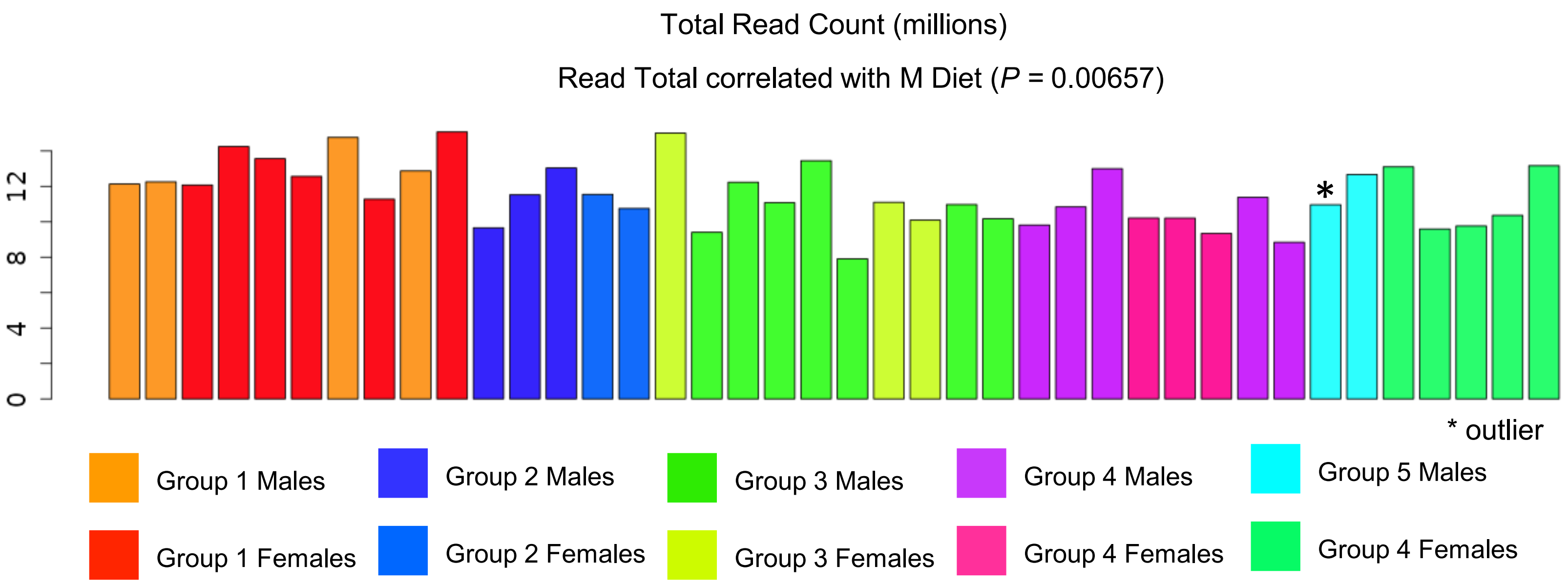

### Figure S3. Impact of maternal diet on gene expression

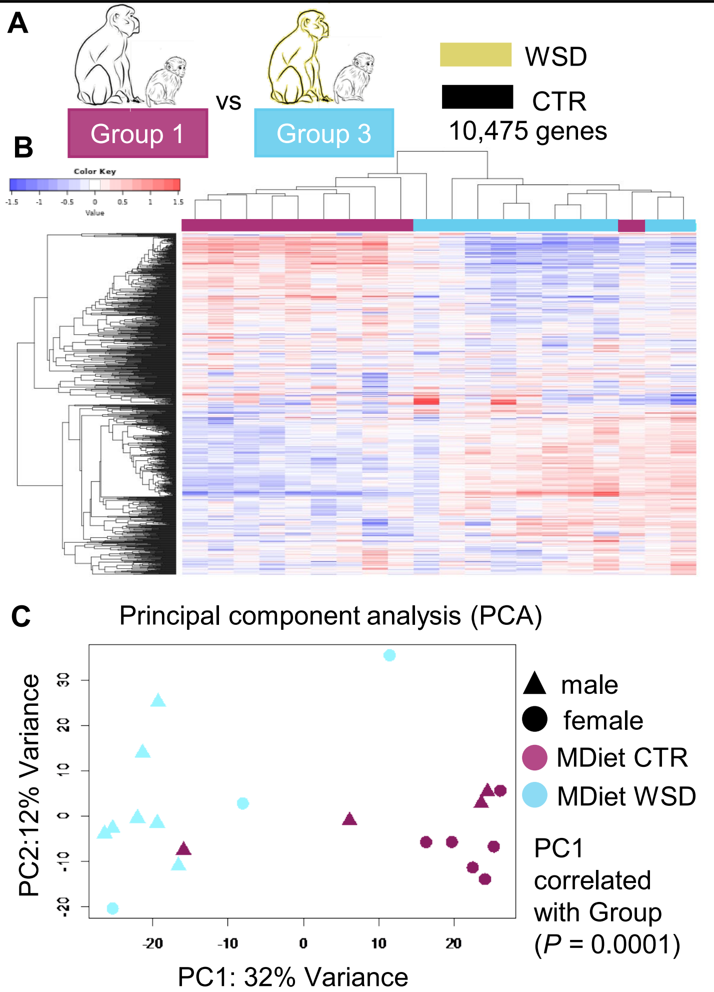


D

Model: ~M Diet

| Comparison | Up | Down |
|---|---|---|
| M Diet WSD vs CTR | **272** | **620** |

### Figure S4. Principal component analysis based on cohort year

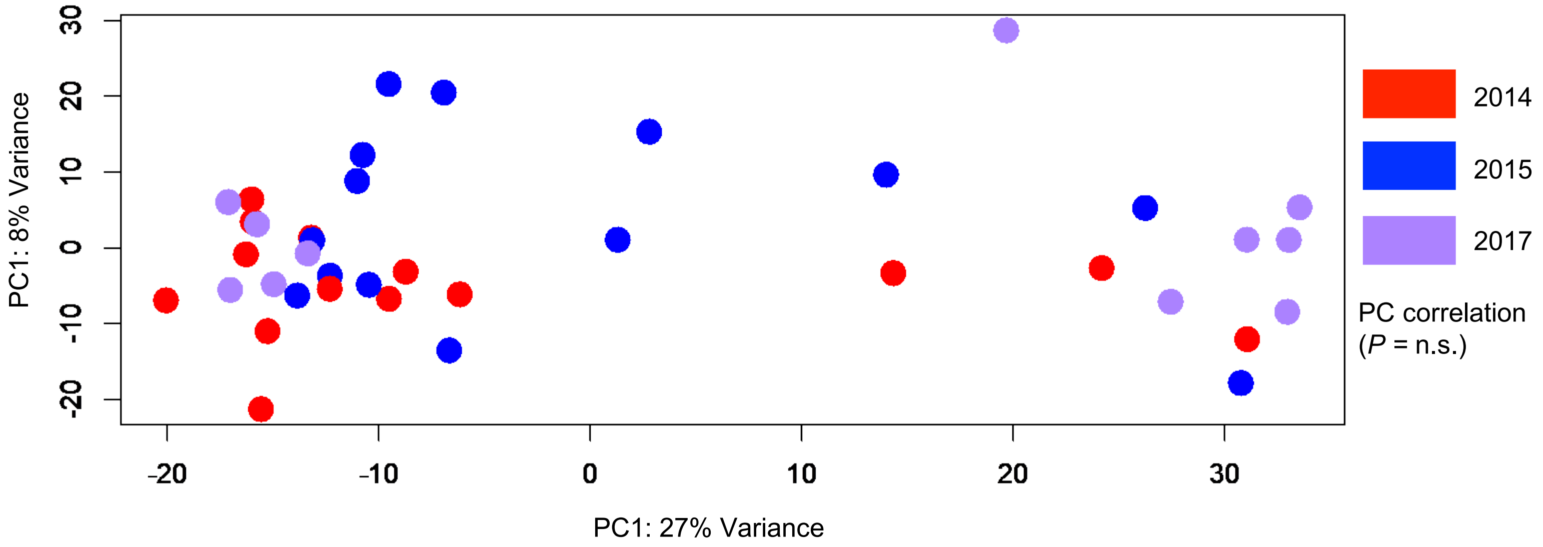

Table S1. Experimental Design Metadata

| Offspring Sample ID | Maternal ID[a] | Offspring Sex | Maternal Age | % Maternal Body Fat[b] | years on WSD | Offspring Collection[c] | Offspring Insulin (µU/mL) | Experimental Group[d] |
|---|---|---|---|---|---|---|---|---|
| T100 | Dam12 | M | 7.3 | 9 | NA | 2014 | 6.1 | Group 1 |
| T101 | Dam10 | M | 9.1 | 17 | NA | 2014 | 4.0 | Group 1 |
| T102 | Dam13 | F | 7.7 | 10 | NA | 2014 | 2.8 | Group 1 |
| T103 | Dam01 | F | 13.1 | 11 | NA | 2014 | 3.3 | Group 1 |
| T104 | Dam18 | F | 8.1 | 24 | NA | 2015 | 18.2 | Group 1 |
| T105 | Dam18 | F | 10 | 24 | NA | 2017 | 12.8 | Group 1 |
| T106 | Dam05 | M | 14 | 21 | NA | 2017 | 4.7 | Group 1 |
| T107 | Dam07 | F | 12.9 | 23 | NA | 2017 | 3.6 | Group 1 |
| T108 | Dam06 | M | 13.1 | 20 | NA | 2017 | 5.2 | Group 1 |
| T109 | Dam09 | F | 12.1 | 25 | NA | 2017 | 6.4 | Group 1 |
| T201 | Dam09 | M | 9 | 13 | NA | 2014 | 30.6 | Group 2 |
| T202 | Dam07 | M | 9.9 | 16 | NA | 2014 | 9.8 | Group 2 |
| T203 | Dam06 | M | 11.1 | 15 | NA | 2015 | 7.4 | Group 2 |
| T204 | Dam05 | F | 12.2 | 15 | NA | 2015 | 18.2 | Group 2 |
| T205 | Dam09 | F | 10.2 | 20 | NA | 2015 | 38.5 | Group 2 |
| T300 | Dam19 | F | 6.2 | 22 | 1.8 | 2014 | 5.2 | Group 3 |
| T304 | Dam14 | M | 7.7 | 11 | 1.2 | 2015 | 3.1 | Group 3 |
| T305 | Dam23 | M | 6 | 9 | 1.2 | 2015 | 3.6 | Group 3 |
| T306 | Dam17 | M | 7.6 | 13 | 1.3 | 2015 | 6.0 | Group 3 |
| T307 | Dam28 | M | 5.2 | 23.3 | 1.4 | 2015 | 5.2 | Group 3 |
| T308 | Dam25 | M | 6 | 16 | 2.6 | 2015 | 1.9 | Group 3 |
| T309 | Dam27 | F | 8.3 | 24 | 4.6 | 2015 | 13.6 | Group 3 |
| T311 | Dam14 | F | 10 | 26 | 3.4 | 2017 | 4.6 | Group 3 |
| T312 | Dam27 | M | 7.2 | 26 | 3.5 | 2017 | 5.1 | Group 3 |
| T313 | Dam22 | M | 8.1 | 27 | 3.5 | 2017 | 4.8 | Group 3 |
| T400 | Dam20 | F | 6.1 | 13.5 | 1.5 | 2014 | 13.1 | Group 4 |
| T401 | Dam11 | F | 7.2 | 12 | 1.7 | 2014 | 3.5 | Group 4 |
| T402 | Dam16 | F | 7.1 | 15 | 1.7 | 2014 | 13.1 | Group 4 |
| T404 | Dam26 | M | 5.1 | 15 | 1.7 | 2014 | 11.4 | Group 4 |
| T405 | Dam22 | M | 6.1 | 9 | 1.6 | 2015 | 9.5 | Group 4 |
| T406 | Dam20 | M | 7.4 | 17 | 2.8 | 2015 | 11.6 | Group 4 |
| T407 | Dam21 | F | 7.8 | 10 | 1.5 | 2015 | 9.5 | Group 4 |
| T408 | Dam11 | F | 8.3 | 10 | 2.7 | 2015 | 7.3 | Group 4 |
| T500 | Dam24 | M | 8.3 | 30 | 4.6 | 2017 | 4.3 | Group 5 |
| T501 | Dam15 | M | 10.2 | 40 | 3.5 | 2017 | 2.2 | Group 5 |
| T502 | Dam02 | F | 12.2 | 35 | 5.8 | 2014 | 4.3 | Group 5 |
| T503 | Dam04 | F | 12.3 | 30 | 5.8 | 2014 | 2.3 | Group 5 |
| T504 | Dam03 | F | 12.4 | 33 | 5.9 | 2014 | 10.1 | Group 5 |
| T505 | Dam08 | F | 12.9 | 35 | 8.5 | 2017 | 10.1 | Group 5 |
| T506 | Dam17 | F | 9.8 | 34 | 3.5 | 2017 | 6.5 | Group 5 |

a- Relatedness analyses were grouped by maternal ID, paternal ID is unknown

b- Body Fat % $\leq$ 27 = Lean, Body Fat % $\geq$ 30% = Obese

c- Rearing cohort determined by year necropsy was performed

Table S2. Adult female characteristics and physiology prior to pregnancy (NP) and during the third trimester (3T)

| | Lean CTR | Lean WSD | Obese WSD |
|---|---|---|---|
| Total offspring/group | 15 | 18 | 7 |
| Number of dams | 9 | 13 | 7 |
| Age at birth | 10.7 ± 0.6 (15) [a] | 7.1 ± 0.3 (18)[b] | 11.0 ± 0.7 (6) |
| Gravid [range] | 4.9 ± 0.6 [3-8] (15) [a] | 2.4 ± 0.3 [1-5] (18)[b] | 5.1 ± 0.8 [2-8] (7) |
| Years on WSD | - | 2.2 ± 0.2(18) * | 5.4 ± 0.7 |
| NP body fat (%) | 17.5 ±1 (15) | 16.6 ± 2 (18) [b] | 33.6 ± 1 (7) [c] |
| NP weight (kg) | 9.0 ± 0.2 (15) | 8.5 ± 0.3 (18) [b] | 12.4 ± 0.4 (7) [c] |
| NP Fasting Insulin (μU/mL) | 14.7 ± 2.9 (13) | 19.8 ± 3.4 (18) | 42.08 ± 18.1 (7) [c] |
| NP Fasting Glucose (mg/dL) | 50.9 ± 2.2 (13) | 53.3 ± 2.9 (18) | 51.1 ± 4.7 (7) |
| NP HOMA-IR | 2.0 ± 0.4 (13) | 2.6 ± 0.4 (18) | 6.4 ± 3.7 (7) |
| NP glucose AUC | 6488 ± 334 (13) | 5582 ± 221 (18) | 6730 ± 607 (7) |
| NP insulin AUC | 2251 ± 325 (13) | 3604 ± 537 (18) | 5405 ± 1006 (7) [c] |
| 3T weight (kg) | 10.0 ± 0.2 (15) | 9.2 ± 0.3 (18) [b] | 12.2 ± 0.7 (7) [c] |
| 3T Fasting Insulin (μU/mL) | 35.8 ± 12.4 (11) | 26.7 ± 6.0 (14) | 50.94 ± 11.6 (7) |
| 3T Fasting Glucose (mg/dL) | 40.6 ± 2.8 (11) | 44.2 ± 3.1 (14) | 38.3 ± 2.7 (7) |
| 3T HOMA-IR | 4.4 ± 1.9 (11) | 3.4 ± 1.1 (14) | 4.6 ± 0.9 (7) |
| 3T glucose AUC | 4650 ± 272 (11) | 3850 ± 279 (13) | 4756 ± 442 (5) |
| 3T insulin AUC | 5491 ± 741 (10) | 5733 ± 667 (13) [b] | 18493 ± 4515 (6) [c] |

Data are means ± SEM. Number of data points included in each measurement is indicated in parenthesis. Glucose and insulin were measured during an i.v. glucose tolerance test and area under the curve (AUC) calculated from baseline. Body fat was measured by DEXA and percent fat calculated from total mass. Data were analyzed by one-way ANOVA with a Tukey multiple comparison test. [a] $P<0.05$ Lean CTR vs. Lean WSD; [b] $P<0.05$ Lean WSD vs. Obese WSD; [c] $P<0.05$ Lean CTR vs. Obese WSD. A Student's t- test was used to compare years on the WSD between lean and obese dams with * $P<0.05$
